# SixPack-AbScan: a web server to discover cross-reactivity of antibodies across species

**DOI:** 10.64898/2026.09.10.746162

**Authors:** Marco Grillo

## Abstract

Availability of commercial antibodies for immunochemistry is typically limited to major model species. Researchers working on non-model organisms therefore often need either to generate novel species-specific antibodies or venture into expensive empirical screening from available antibody catalogues, hoping to find cross-reactive reagents. SixPack-AbScan is a free web server aiding researchers in transferring antibodies across species: using the available epitope-mapping information, the software performs a simple computational pre-screening of potential cross-reactivity. The workflow is species-agnostic and designed to help non-model-species researchers prioritize antibodies for experimental validation. A hit indicates sequence-level conservation of the epitopes and a high probability of cross-reactivity; the server does not model substitutions, structure, accessibility, expression or binding affinity. The web server is available at https://sixpack-abscan.serve.scilifelab.se.

**GRAPHICAL ABSTRACT:** 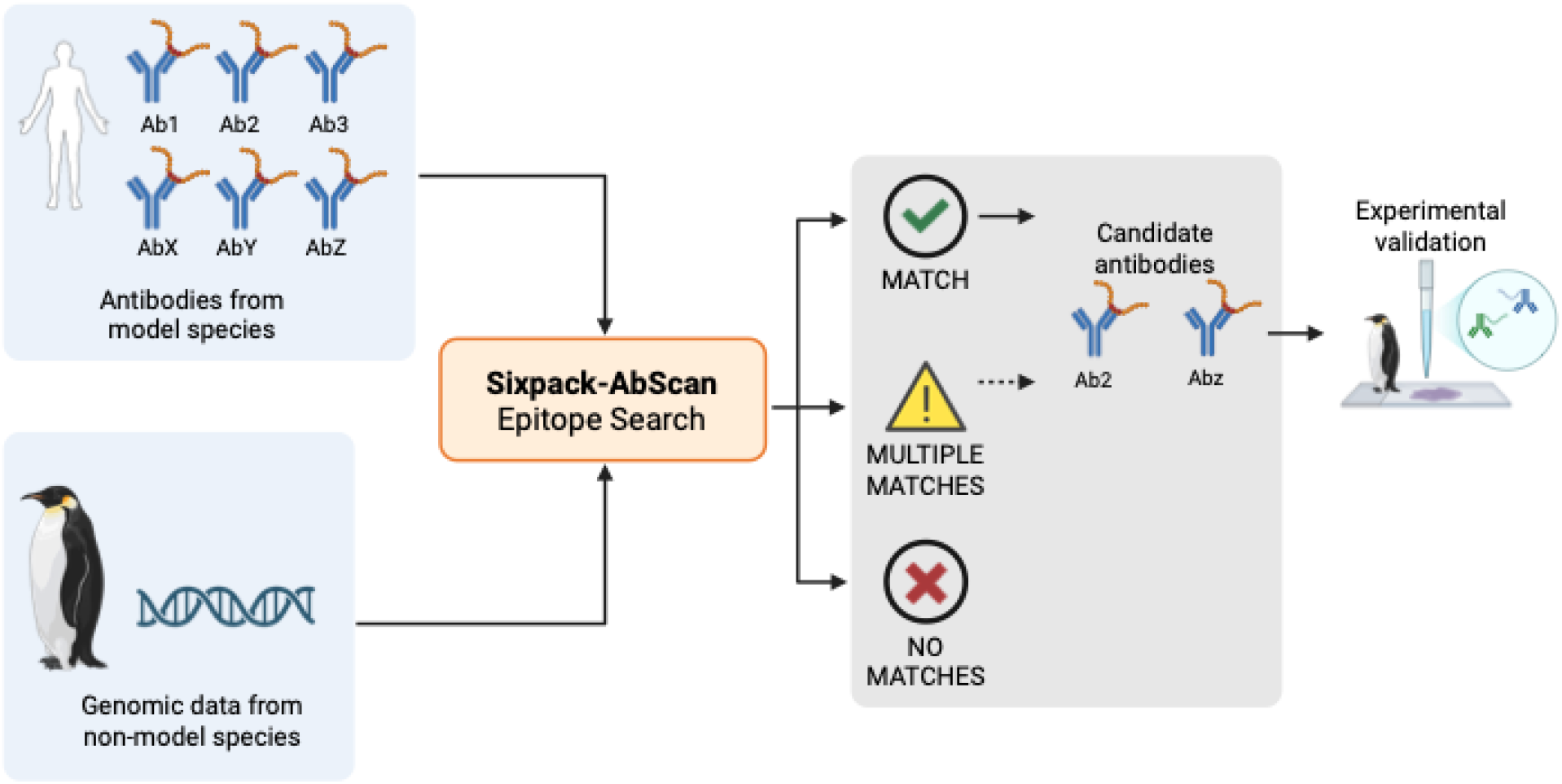

## INTRODUCTION

Research antibodies are widely used in immunochemical applications, including immunohistochemistry and Western blotting (1). However, most commercially available antibodies have been developed for a relatively small group of species (2). These include organisms of direct clinical relevance, such as humans, mice, and rats, as well as experimentally tractable model organisms supported by large research communities, such as *Drosophila, Caenorhabditis elegans*, zebrafish, and *Arabidopsis*.

Expanding antibody availability beyond this restricted set of species would greatly facilitate studies of biological diversity and non-model organisms (2). Commercial support for such species, however, remains limited. Researchers can always generate antibodies specifically against proteins from their organism of interest, but antibody production is generally expensive and time-consuming. Alternatively, researchers may attempt to repurpose antibodies developed for model organisms, by experimentally screening large collections against a non-model species. For example, antibodies raised against mouse proteins may be tested individually on appropriate chicken tissue and scored for reactivity. This approach has been widely used: when an antibody produces a specific signal in the non-target species, it is therefore described as **cross-reactive** and generally adopted in the new species. Beautiful and extensive use of this approach can be found, for example, in the seminal book “Evolution of Nervous Systems” (3).

Research antibodies can be broadly divided into monoclonal and polyclonal types. Monoclonal antibodies consist of a single antibody species that binds to a defined region of the target antigen. Polyclonal antibodies are mixtures of antibodies that typically recognize several distinct regions of the same antigenic protein.

Monoclonal antibodies offer several advantages: first, they can be maintained and produced from hybridoma cell lines, allowing the same reagent to be manufactured across multiple batches and over long periods of time, guaranteeing long-term experimental reproducibility (1).

Second, monoclonal antibodies can be **epitope-mapped** via biochemical assays that can identify the specific antigenic region (epitope) bound by the antibody (4–6). For antibodies recognizing linear epitopes, peptide-scanning approaches can often localize binding to a short contiguous region of the antigen, in the size range of 10-20 aminoacids. Deletion and substitution analyses within this region can then be used to identify the minimal epitope and the residues that are critical for binding (4).

Once the epitope for a monoclonal antibody is known, its potential reactivity in a non-target species can, in principle, be predicted computationally. Two basic questions can be asked: i) is the recognized amino-acid sequence present in the proteome of the non-target species, and ii) how many proteins might contain that sequence? The presence of an identical peptide in a candidate protein provides a conservative indication that cross-reactivity may be possible. The number and identity of matching proteins can also help anticipate whether the antibody is likely to produce a unique signal, or an ambiguous one (i.e. if the antibody is predicted to recognize multiple proteins in the species of interest).

Several commercial suppliers now maintain substantial collections of epitope-mapped monoclonal antibodies, and the corresponding epitope sequences are often publicly available or can be obtained from the manufacturers upon request. It is therefore possible to computationally pre-screen these antibody collections for cross-reactivity against any given non-model organism, provided that a predicted proteome is available.

Whenever a predicted proteome is not available, a 6-frame translation of the transcriptome, or even of the genome assembly, can be used instead. Translation of the complete nucleotide assembly in all six reading frames generates a large number of putative amino-acid sequences that do not exist but the probability that a specific, high-complexity peptide of this length occurs by chance in an unrelated six-frame translation is extremely low, even when the entire genome is searched. Accidental matches may be more frequent for shorter, repetitive or compositionally biased epitopes, which should therefore be flagged or excluded from interpretation. However, these epitopes are unlikely to exist in the real antibody collections, as they would yield poor specificity in their binding.

Despite the relative conceptual simplicity of this analysis, this workflow may be inaccessible to researchers who are unfamiliar with sequence analysis of nucleotide assemblies, with the inner workings of monoclonal epitope-mapping procedures, or with proteome-wide peptide searches. I aim to address this specific gap, presenting a practical framework for systematic transfer of epitope-mapped monoclonal antibodies to non-model organisms. By integrating mapped antibody epitopes with proteome or six-frame translated sequence searches, SixPack-AbScan enables rapid computational pre-screening of candidate cross-reactive antibodies prior to experimental validation, in a user-friendly format.

Here, we describe the underlying algorithm, web implementation, output schema, intended applications in non-model organisms, and the limitations that require subsequent experimental validation.

## MATERIALS AND METHODS

### Software availability and architecture

The public web interface is hosted by SciLifeLab Serve at https://sixpack-abscan.serve.scilifelab.se and is intended to be accessible over HTTPS without registration. Source code is available under the MIT License at https://github.com/mgcizzu/SixPack-AbScan. A container image is built from a Python 3.11 base image through an automated GitHub Actions workflow and published to GitHub Container Registry.

#### Access

web server | source code | persistent application record

The computational core is implemented in Python. FASTA parsing uses Biopython SeqIO (7), tabular processing uses pandas (8), Excel input uses openpyxl (9), and the interactive interface is implemented with Gradio (10). The same core workflow is also available through a command-line entry point for reproducible local use. The container runs as an unprivileged user and writes run products to application-owned directories.

### Inputs and epitope preprocessing

Each run requires one sequence file and one epitope table. The sequence input can be either a nucleotide FASTA, as genomic or transcriptomic sequence, or a precomputed protein FASTA.The files can be supplied as fasta (either uncompressed or compressed in .gz), or as direct links to NCBI-hosted files, in which cases they get downloaded directly in the app and processed. The epitope table can be supplied as CSV, TSV, TXT, XLSX or XLS. For delimited files the user specifies the separator; for all table types the interface reads the header and allows the peptide-containing column to be selected.

Peptide values are converted to strings, stripped of leading and trailing whitespace, converted to uppercase and stripped of internal space characters. Empty values are removed and the remaining peptides are deduplicated before scanning. Original rows are retained separately so that all antibody or reagent metadata can be restored after the search. The present implementation does not reject non-standard amino-acid symbols; users should therefore inspect input quality before interpreting results.

### Six-frame translation

For nucleotide input, each FASTA record is converted into six amino-acid sequences: offsets 0, 1 and 2 on the submitted strand, followed by offsets 0, 1 and 2 on its reverse complement. Translation uses by default the standard genetic code, but alternative codes can be also specified through a drop-down menu. Stop codons are represented by an underscore and unrecognized codons by X. Reverse complementation handles A, C, G, T and N explicitly; other ambiguity codes propagate into unrecognized codons. Output identifiers append |frame1 through |frame6 to the source record identifier. When providing genomic sequences, this approach might produce unrealistic aminoacid stretches: the 6-frame translation is enforced on the entire genome, on both coding and non-coding regions, likely producing many translations that do not actually exist in the real proteome. However, the chance of these random artifacts to contain by accident a specific non-random 10-20 aminoacid epitope is very low.

### Exact epitope scanning and output generation

The selected protein FASTA, either uploaded or generated via 6-frame translation, is parsed record by record. For each record, each epitope sequence is searched as a literal contiguous substring. A positive result produces one row containing epitope_query, target_id and target_description. The current algorithm records presence per epitope-target pair; it does not report match position, the number of repeated occurrences within a record or local alignment.

Two CSV outputs are created. epitope_hits.csv contains the unique query-target pairs returned by the scan. matched_epitope_rows.csv joins those hits back to every non-empty source row carrying the same normalized peptide, thereby preserving product numbers, gene identifiers and other submitted columns. When a nucleotide input is used, output6frame.fasta is also available. Results are displayed in the browser and can be downloaded individually.

### Private catalogue mode

It is understandable that some companies might want to keep the precise epitope mapping information confidential. For this eventuality, the web server implements a “Private catalogue mode”. This modality allows antibody manufacturers to share the mapping information with the author or the admins of the web server through private communication channels (and under non-disclosure agreements, if required). The mapping information is then pre-loaded in the app, loaded by the user through a dropdown menu, the search is performed in the backend, but the output never exposes nor retains the scanned and matched epitope sequence. In this mode, however, the user is still able to determine whether a given antibody is predicted to cross-react and download the rest of the metadata. I believe this to be an acceptable tradeoff for most manufacturers. As a real example, the web app already hosts a subset of the DSHB catalogue (Developmental Studies Hybridoma Bank), specifically a set of antibodies whose epitope-mapping information was publicly available through the DSHB web page and easy to parse. More catalogues will be added in the future, as they become available.

### Web execution and temporary data

The Gradio interface reports the progress of each step: upload, translation and search progress, and displays both result tables before download. Outputs are written under timestamped run directories. Directories created by a server process are removed on normal process exit, and run directories left by an earlier process are removed at application start.

## RESULTS

### Interactive analysis workflow

A complete run (Figure 1) requires the user to provide three inputs: a table containing at least a list of epitopes recognized by each antibody, or choose a pre-loaded manufacturer input table (Figure 2), the sequence-input mode (protein or nucleotide sequences, as menu choice) and the FASTA file (protein or nucleotide sequence) to scan (Figure 3). Choosing nucleotide input adds a 6-frame translation step, whereas protein input proceeds directly to the search. Progress messages report the number of source sequences, peptide records, translated records, scanned unique epitopes and accumulated hits. On completion, the interface displays the hit table and metadata-enriched table and exposes the generated files for download (Figure 4).

**Figure 1.**
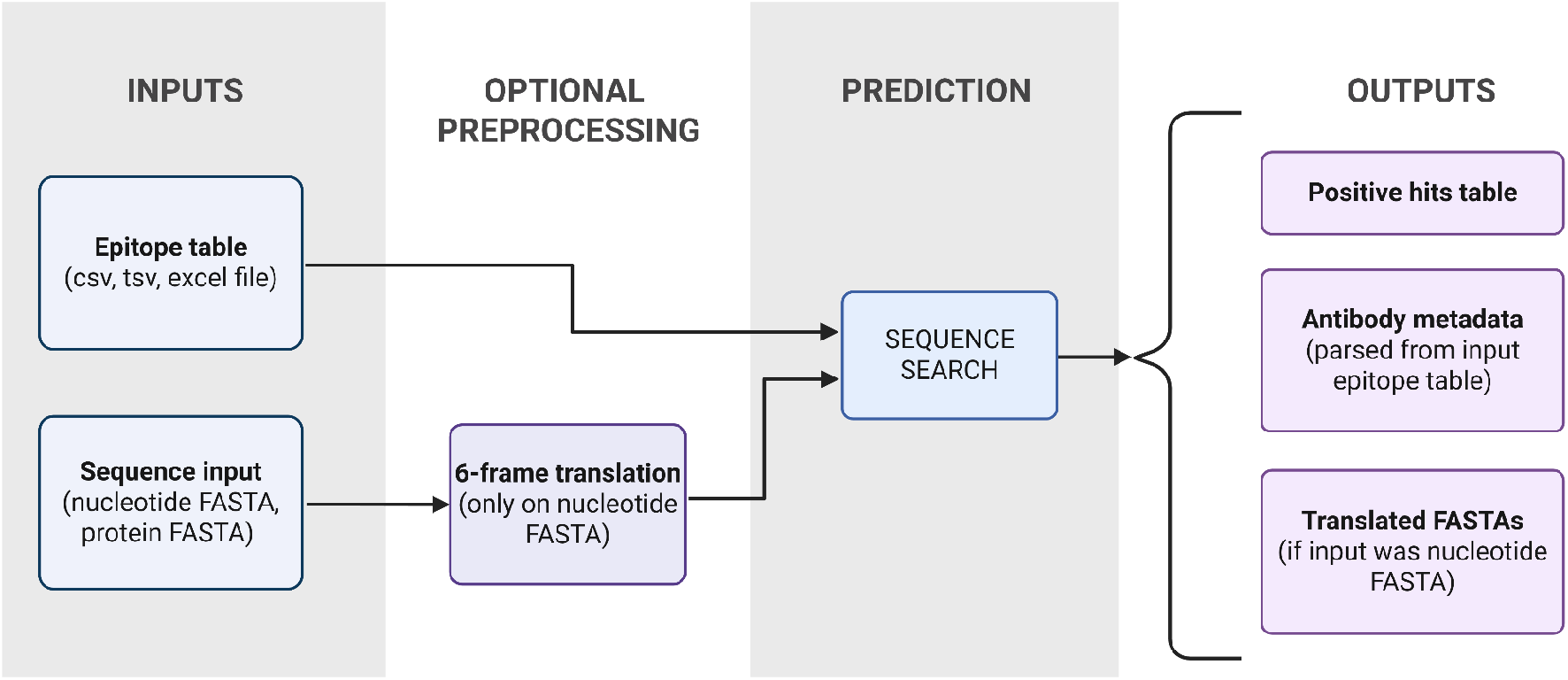
SixPack-AbScan workflow. A table of mapped peptide epitopes is screened against either a submitted proteome or all six translations of a nucleotide FASTA. Translation can be executed according to all the NCBI translation codes. The server reports matching target records and restores the source antibody metadata. Exact sequence identity is interpreted as a compatibility flag for non-model-species antibody transfer, not as proof of antibody binding.

**Figure 2.**
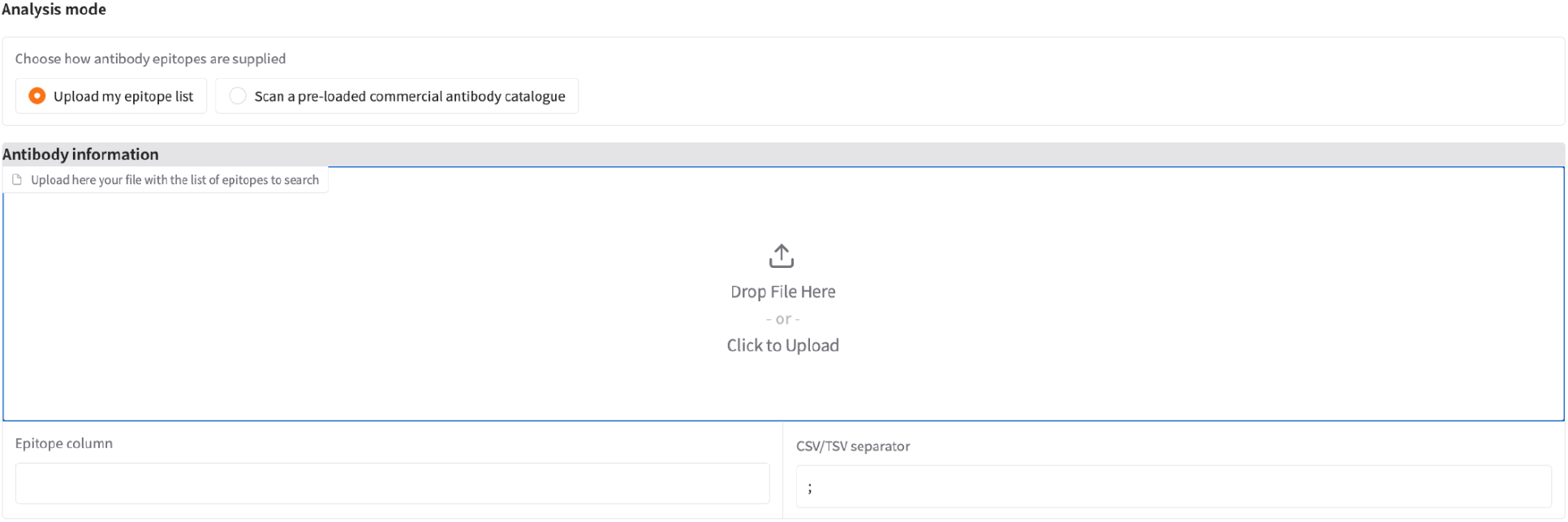
Input window for the Antibody information: A table of mapped epitopes needs to be uploaded by the user. The user needs to specify the header of the Epitope column (the column containing the epitope sequences), as well as a separator type (if the file is CSV/TSV). The headers of all the columns will become available as a dropdown list upon uploading of a file. In the case of an Excel table, the separator information does not need to be specified.

**Figure 3.**
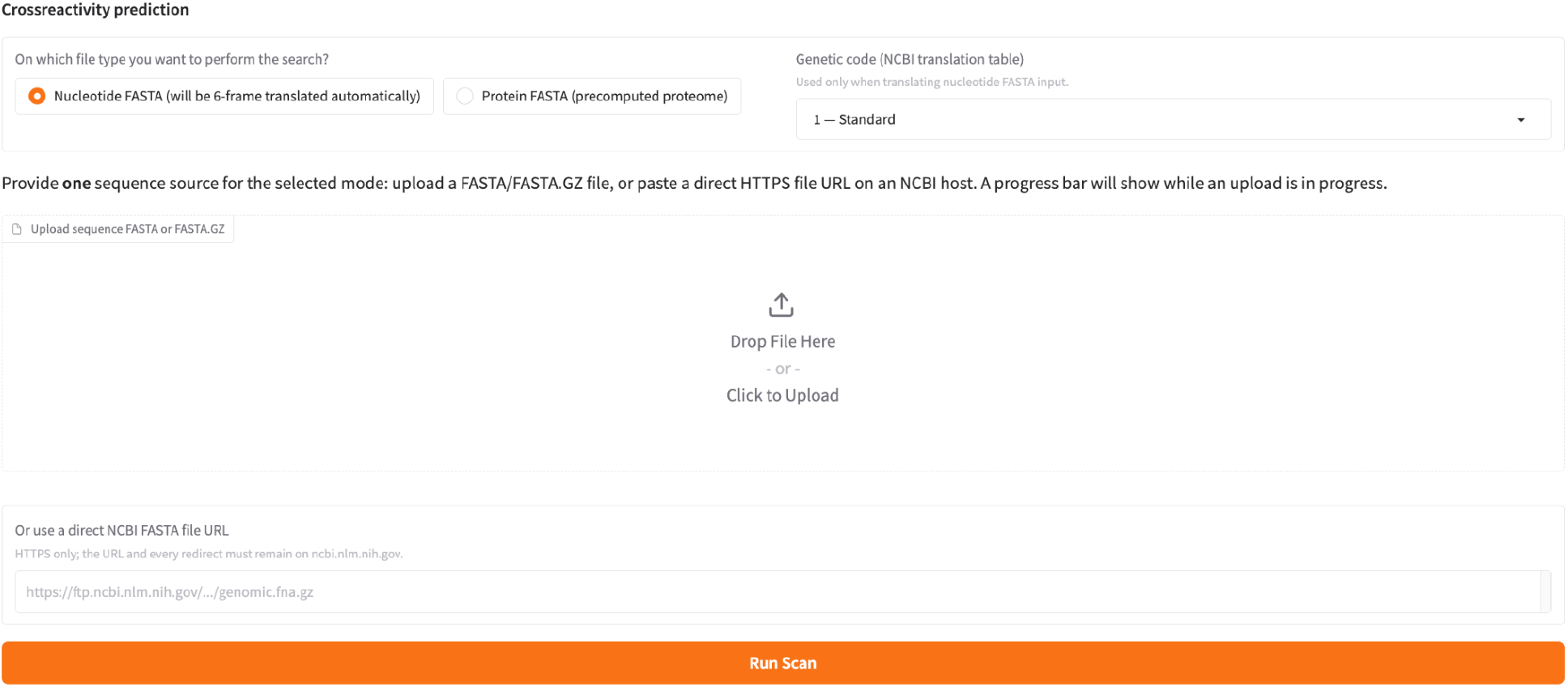
Input window for the target proteome information: A FASTA file needs to be uploaded by the user, the file can be in .fasta or fasta.gz format, or can be directly input as a NCBI link. The user needs to specify if the input file is a nucleotide or protein sequence. If the input file is a nucleotide, it will be 6-frame translated, and the translations will be scanned for the epitope sequences. The app supports the choice of alternative genetic codes for translation, via a dropdown menu.

**Figure 4.**
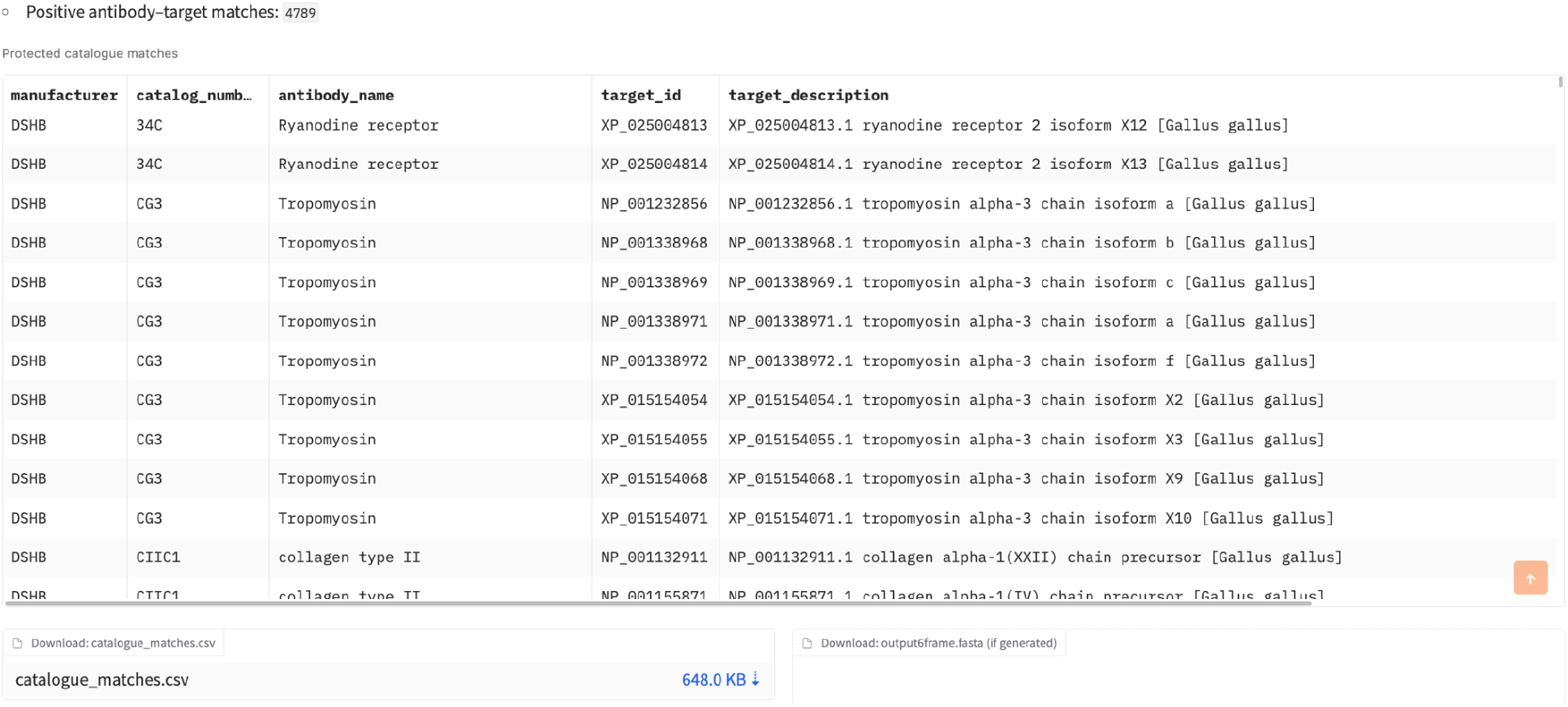
Example output window for a proteome run: A list of positive matches epitopes is reported, specifying the positive hit identity on the target proteome (top panel). For each positive match, a full list of the antibody, the target protein and other metadata (such in this case the catalogue number) are reported on screen. If the 6-frame translation is perform can be downloaded as .csv files.

### Example input profile

The bundled workbook, loaded through the “Scan a pre-loaded commercial antibody catalogue” illustrates a real case use with repeated antibody products, as well as shared epitopes between multiple antibodies (product and epitope list retrieved from the DSHB product pages). It contains product rows spanning distinct protein symbols, Ensembl identifiers and UniProt identifiers (these data refers to the proteins targeted by the antibodies in the original target species).

### Intended use

The primary use case is early reagent triage: an exact hit in a new species indicates potential cross-reactivity and supports prioritizing that antibody for experimental testing. Multiple search hits support the existence, in the target species, of multiple proteins carrying the epitope and potentially recognized by the antibody (i.e. closely related proteins, or highly conserved domains). In other words: a specific antibody from species A could potentially bind more than one target protein from species B. Because the output keeps the normalized peptide, target FASTA context and every original metadata row, these decisions can be traced to a specific commercial product and reviewed by the user.

These interpretations are intentionally conservative. SixPack-AbScan does not establish orthology, expression, epitope accessibility, native folding or affinity; the user must inspect the target identifiers and validate the selected reagent in the intended assay and species.

### Relationship to existing web resources

SixPack-AbScan can screen any user-supplied species and antibody list. This positioning makes the server most appropriate for transparent, early-stage triage of commercial antibodies in organisms for which antibodies, or even a curated proteome dataset, may be unavailable. To my knowledge, no other web resource exists allowing a similar exploration.

### Interpretation of results

A reported hit for an antibody means that one or more identical epitopes occur in a submitted protein record or translated reading frame. Absence of a hit is not evidence that an antibody will fail to bind, because conservative substitutions, short motifs, conformational mimicry and incomplete assemblies are not contemplated by the search. Likewise, a hit does not guarantee that the antibody will work, as this also depends on expression, accessibility, or post-translational modifications of the epitope.

## DISCUSSION

SixPack-AbScan converts a recurring practical question into a reproducible and user-friendly web workflow: can commercial antibodies used in established model organisms also be used on a non-model organism with a reasonable chance of success?

The contribution of SixPack-AbScan is primarily integration and ease of use. Six-frame translation removes the need to prepare a predicted proteome when only a genome or transcriptome assembly is available, batch tables allow many reagents to be evaluated in one run, and the output results are returned in a form that can be traced to product numbers and target identifiers. The literal matching rule is conservative, and intuitively useful to prioritize experiments.

The conservative design also imposes substantial limitations. Only continuous linear epitopes are represented. The program does not score substitutions, gaps or biochemical similarity; it does not consider solvent exposure, secondary or tertiary structure, post-translational modification, expression or antibody affinity. Six-frame translation does not predict open reading frames or require start codons, so matches in theory can arise in translated non-coding sequences. However, the probability of this actually happening for a complex epitope is extremely low. Assembly errors, contamination and incomplete sequence coverage can in theory generate false candidates or missed matches.

These constraints define the intended use. SixPack-AbScan should be treated as a pre-screening tool for non-model-species antibody transfer: it identifies exact sequence conservation and obvious sequence-level targets, after which candidate reagents still require experimental validation in the intended assay and species. Suitable follow-ups may include the integration of experimental data, whenever available.

In summary, SixPack-AbScan provides a species-independent rapid, interpretable triage of potentially cross-reactive antibodies, targeted to researchers working on non-model species.

## DATA AVAILABILITY

SixPack-AbScan is available without login at https://sixpack-abscan.serve.scilifelab.se. Source code is available at https://github.com/mgcizzu/SixPack-AbScan under the MIT License. The current deployed application record is https://doi.org/10.82595/scilifelab.4c3a-tp57 and the all-versions record is https://doi.org/10.82595/scilifelab.34k8e-6fr80.

## ACKNOWLEDGEMENTS

The authors acknowledge the SciLifeLab Serve hosting infrastructure, Michalis Averof for the original insight on the epitope search, Mats Nilsson for useful suggestions on the manuscript and the Developmental Studies Hybridoma Bank (DSHB) for openly sharing the epitope-mapping information on their product pages.

## AUTHOR CONTRIBUTIONS

Marco Grillo: Conceptualization, Methodology, Software, Validation, Visualization, Writing—original draft, Writing—review and editing. Revise the CRediT roles if additional authors or contributors are added.

## REFERENCES

1. Uhlen, M., Bandrowski, A., Carr, S., Edwards, A., Ellenberg, J., Lundberg, E., Rimm, D.L., Rodriguez, H., Hiltke, T., Snyder, M., et al. (2016) A proposal for validation of antibodies. Nat. Methods, 13, 823–827.

2. Dhar, P., Samarasinghe, R.M. and Shigdar, S. (2020) Antibodies, nanobodies, or aptamers-which is best for deciphering the proteomes of non-model species? Int. J. Mol. Sci., 21, E2485.

3. Kaas, J.H., Striedter, G.F., Bullock, T.H., Preuss, T.M., Rubenstein, J.L.R. and Krubitzer, L.A. eds (2017) Evolution of Nervous Systems 2nd edn Academic Press, Oxford.

4. Reimer, U., Reineke, U. and Schutkowski, M. (2011) Peptide arrays for the analysis of antibody Epitope recognition patterns. Mini Rev. Org. Chem., 8, 137–146.

5. Buus, S., Rockberg, J., Forsström, B., Nilsson, P., Uhlen, M. and Schafer-Nielsen, C. (2012) High-resolution mapping of linear antibody epitopes using ultrahigh-density peptide microarrays. Mol. Cell. Proteomics, 11, 1790–1800.

6. Michaud, G.A., Salcius, M., Zhou, F., Bangham, R., Bonin, J., Guo, H., Snyder, M., Predki, P.F. and Schweitzer, B.I. (2003) Analyzing antibody specificity with whole proteome microarrays. Nat. Biotechnol., 21, 1509–1512.

7. Cock, P.J.A., Antao, T., Chang, J.T., Chapman, B.A., Cox, C.J., Dalke, A., Friedberg, I., Hamelryck, T., Kauff, F., Wilczynski, B., et al. (2009) Biopython: freely available Python tools for computational molecular biology and bioinformatics. Bioinformatics, 25, 1422–1423.

8. The pandas development team (2026) pandas-dev/pandas: Pandas Zenodo.

9. Clark, S.R.F.C. (2026) openpyxl: A Python library to read/write Excel 2010 xlsx/xlsm files.

10. Abid, A. and Others (2019) Gradio: Hassle-Free Sharing and Testing of ML Models in the Wild. arXiv preprint arXiv:1906. 02569.

